# Discovering cancer driver genes and pathways using stochastic block model graph neural networks

**DOI:** 10.1101/2021.06.29.450342

**Authors:** Viola Fanfani, Ramon Vinas Torne, Pietro Lio’, Giovanni Stracquadanio

## Abstract

The identification of genes and pathways responsible for the transformation of normal cells into malignant ones represents a pivotal step to understand the aetiology of cancer, to characterise progression and relapse, and to ultimately design targeted therapies. The advent of high-throughput omic technologies has enabled the discovery of a significant number of cancer driver genes, but recent genomic studies have shown these to be only necessary but not sufficient to trigger tumorigenesis. Since most biological processes are the results of the interaction of multiple genes, it is then conceivable that tumorigenesis is likely the result of the action of networks of cancer driver and non-driver genes.

Here we take advantage of recent advances in graph neural networks, combined with well established statistical models of network structure, to build a new model, called Stochastic Block Model Graph Neural Network (SBM-GNN), which predicts cancer driver genes and cancer mediating pathways directly from high-throughput omic experiments. Experimental analysis of synthetic datasets showed that our model can correctly predict genes associated with cancer and recover relevant pathways, while outperforming other state-of-the-art methods.

Finally, we used SBM-GNN to perform a pan-cancer analysis, where we found genes and pathways directly involved in the hallmarks of cancer controlling genome stability, apoptosis, immune response, and metabolism.

## 1 Introduction

The classical paradigm of cancer formation suggests that tumors arise from the stochastic accumulation of somatic mutations in key genes, called cancer driver genes, which give aberrant cells the ability to escape cell death and immune response and to grow uncontrollably throughout the body [1, 2].

The advent of high-throughput sequencing technologies has enabled the study of a broad spectrum of cancers and the identification of hundreds of driver genes across different malignancies [3, 4]. While the causal role of many genes in cancer has been confirmed by in-vitro and in-vivo models, recent studies have shown that driver mutations occur also in normal tissues [5, 6]. These observations suggest that driver mutations are necessary but not sufficient for tumorigenesis, and that the order in which they are acquired and the joint alteration of non cancer driver genes is important to transform a normal cell into a malignant one.

While the evolution of cancer cells can now be studied at sufficient resolution to generate testable hypotheses, discovering pathways of driver and non-driver genes associated with cancer has been challenging [7]. Current high-throughput omic assays provide only information with single-gene resolution, whereas gene and protein interaction experiments provide only pairwise information. It has now become apparent that methods able to perform multi-omic analyses at the pathway level are pivotal to understand the aetiology of cancer and design effective therapies.

In the last ten years, there have been substantial efforts to develop network analysis methods that would capture cancer poligenicity [7]. Nonetheless, current approaches typically focus on predicting either new cancer driver genes [8] or cancer driving pathways [9] by integrating multi-omic information; however, these methods usually aggregate multiple experimental information into a gene-level score, which effectively masks the effects and relationships between multiple biological processes underpinning cancer phenotypes.

Here we addressed current limitations in network-aware cancer analysis by developing a new method to simultaneously discover cancer driver genes and pathways by integrating gene-level high-throughput experiments with protein interaction information. To do that, we built a new deep learning model, combining graph neural networks (GNNs, [10]) and stochastic block models (SBMs, [11]), called Stochastic Block Model Graph Neural Network (SBM-GNN). GNNs provide a framework to obtain network-aware embeddings of gene level features; these models have been successfully applied to a number of tasks, including node labelling and link prediction [12, 10, 13], and have been shown to provide meaningful representations of omic data [14]. Embeddings are then combined with SBMs, a robust generative framework to model network connectivity, to infer new pathways. Importantly, our model can be fit end-to-end using standard gradient descent and scales efficiently with the size of the datasets.

To assess the performances of our method for discovering cancer driver genes and pathways, we built a simulation framework to generate synthetic networks with different structures and feature-level multi-modalities; experimental results show that our method is able to detect cancer driver genes and pathways with high accuracy. We then applied SBM-GNN to pan-cancer genome data [3]; here we found that our method outperforms other state-of-the-art approaches in identifying cancer driver genes, while being able to discover pathways associated with the hallmarks of cancer.

## 2 Methods

### 2.1 Model architecture

Let 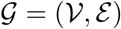 be an undirected graph, with 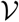 being the set of *n* vertices (or nodes) representing genes or proteins, and *ε* being the set of *m* edges (or links) between nodes in 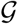. Edges can be represented by an adjacency matrix **A** ∈ ℝ^*n×n*^, such that **A**_*uv*_ = 1 iff node *u* is connected to node *v* in 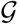, and 0 otherwise. We define **X** ∈ ℝ^*n×f*^ as the matrix of *f* gene features (e.g. mRNA abundance, number of somatic mutations), and **Y** ∈ {0, 1}^*n×c*^ a matrix of *c* gene labels (e.g. being a cancer driver gene), such that **Y**_*uq*_ = 1 iff node *u* has label *q* and 0 otherwise.

Here we hypothesise that node labels depend on the observed gene features and their involvement in pathways mediating the phenotypes of interest; this information is obviously unknown but can be learned from the data.

We denote with Ŷ, 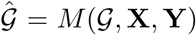 a model that takes in input a graph 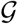, a feature matrix **X**, and a label matrix **Y** and predicts new labels Ŷ and a new graph 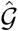. Our model consists of three layers: a network-aware embedding layer, a community detection layer, and a label prediction layer.

The network-aware embedding layer is used to learn a latent dense representation of the biological processes mediated by each gene and its immediate neighbours. The embedding for **X** can be computed using a non-linear transformation, *ψ*, defined as:

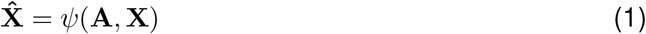

where 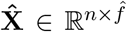 is an 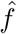-dimensional embedding conditioned on the observed node features **X** and the adjacency matrix **A** of the graph 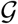.

We then wanted to detect how genes are organised in pathways. To do that, we implemented a community detection layer, which allows us to identify groups of nodes that are strongly connected to each other and that have homogeneous features. To perform community detection, we used a layer that models network structure using Stochastic Block Models (SBM) [15]. SBM is a generative model where edges between nodes depend on a community matrix **B** ∈ [0, 1]^*k×k*^ and a membership matrix **Z** ∈ [0, 1]^*n×k*^, where *k* is the number of unknown blocks. Each entry of the membership matrix, **Z**_*ik*_ denotes the probability that a node *i* belongs to the *k*-th block, which effectively corresponds to assigning genes to pathways. In a canonical SBM, **Z** is binary; however, genes often belong to multiple pathways, thus we relaxed this constraint by allowing for mixed membership, s.t. Σ_*k*_ **Z**_*ik*_ = 1. Each entry of the community matrix **B**_*ij*_, instead, denotes the probability of observing an edge between two nodes belonging to blocks *i* and *j*, respectively (see Supplementary Materials, Supplementary Figure 1). In practice, we infer **B** from the matrix of observed edges **C** = **ZAZ**^*T*^, where **C**_*ij*_ is the number of edges between blocks *i* and *j* [16].

However, since cancer is not only mediated by short-range but also long-range interactions, a naive approach combining a network embedding layer with a SBM layer will likely lead to poor performances. To overcome this problem, we designed our model to simultaneously learn multiple SBMs with a decreasing number of blocks, as a way to force the model to capture both short and long-range interactions. Here we defined a multi SBM layer, as a layer consisting of *S* SBMs with different numbers of blocks, *k_s_*. W.l.o.g. we assume *s* = 0 to index the SBM with the smallest number of blocks; intuitively, high index SBMs represent fine-grained communities, whereas low index SBMs represent coarse-grained communities.

To learn the membership matrix **Z**^(*s*)^ for the *s*-th SBM, we apply a non linear network-aware transformation *ζ* to the embedding of the node features, 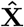, as follows:

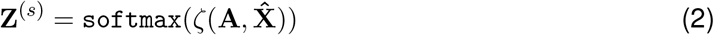

where the `softmax` transformation ensures that 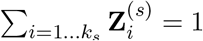.

Finally, to perform node label prediction, we concatenated *S* membership matrices **Z**^(*s*)^ column-wise, and use the resulting matrix as input for the output layer, *ϕ*, as follows:

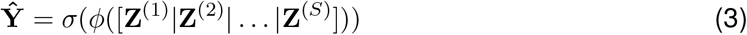

where *σ* is a non-linear transformation suitable for binary or multi category classification.

#### 2.1.1 Learning parameters of an SBM-GNN model

To fit our model, we defined a differentiable loss function, 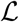, as follows:

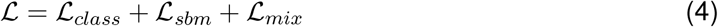

where 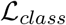 is a supervised loss function for label prediction, whereas 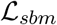 and 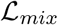 are unsupervised loss functions for learning communities. Specifically, given a membership matrix **Z** and a block matrix **B**, 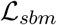 is the likelihood of observing *m* edges in 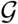 defined as:

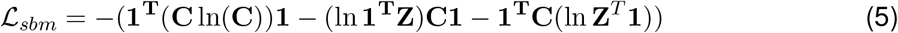

and which in turn is averaged across each SBM layer [16]. Moreover, the contribution of 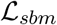 is weighted as a function of the number of epochs, such that the learning process is forced to minimise the community loss first and to learn how to classify the nodes later.

However, fitting multiple SBMs tends to assign nodes to only few blocks, effectively skipping learning community structures. To overcome this problem, we introduced a membership loss function, 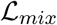, defined as:

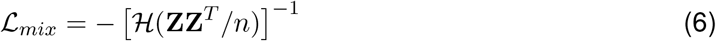

where 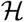 is the entropy function and *n* is the number of nodes; in practice, 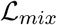 penalizes model configurations assigning all nodes to a single community.

#### 2.1.2 Implementation

Our architecture can be easily tailored to different type of data and analyses by using appropriate transformations for *ψ, ζ, ϕ*. Our goal is to predict cancer driver genes and cancer associated pathways; thus, we defined our base model as follows:

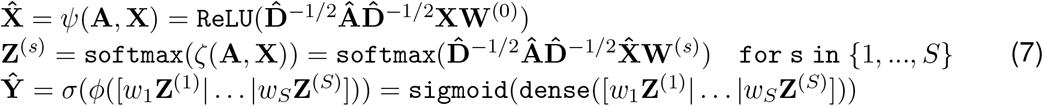

where 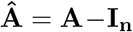 is the and 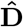 is the node degree matrix of 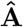. Layers *ψ* and *ζ* are implemented using Graph Convolutional Network (GCNs, [10]) layers, and *ϕ* is a fully connected layer. Importantly, we rescale membership information by learning weights, *w_s_*, in order to identify the most relevant blocks with respect to the node labelling task.

We also explored other architectures, where we changed the *ψ* and *ζ* layers, while keeping fixed the fully connected layer to perform label prediction; we denoted each model configuration using a positional notation (see Table 1).

**Table 1:** SBM-GNN implementations. For each implementation used in our study, we report the type of layer used for the network-aware embedding function, *ψ*, and the membership assignment function, *ζ*.

| Model | $\psi$ | $\zeta$ |
| --- | --- | --- |
| GCN-GCN | GCN | GCN |
| LIN-GCN | fully connected | GCN |
| GCN2-GCN | GCN depth 2 | GCN |
| GCN-LIN | GCN | fully connected |

A possible limitation of our base model is that gene labels might be strongly correlated to gene features, and this information might not be captured only by the SBM layers. Here, we addressed this limitation by concatenating the embedding of gene-level features to the membership vectors, such that 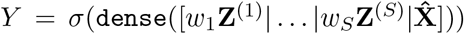; we refer to this layer as a residual layer (denoted by the suffix RES in the name) [17].

We also considered introducing pre-processing steps to speed up learning model parameters by augmenting our input node features. Biological network analyses have shown that graph diffusion is a powerful technique to extract information from protein interaction data ([18]). Thus, we decided to also test the use of graph diffusion as a pre-processing step using a sparsified version of a random walk diffusion matrix, called Graph Diffusion Convolution layer (GDC, [19], denoted by the prefix GDC in the model name).

Taken together, we tested 4 different model architectures that we then trained with either pre-processed and raw node features, and using residuals (see Table 1).

### 2.2 Synthetic data simulation

#### 2.2.1 Synthetic gene networks

We simulated networks as a mixture of a perfect communities graph 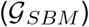 and a random Erdos-Renyi network 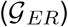 [20]. By varying a noise parameter *η*, we parametrised the contribution of noise and planted communities as 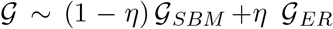, whereas network sparsity is controlled by a density factor *d* (see Supplementary Materials, Supplementary Figure 2). As already shown, a stochastic block model with *k* communities and *n* nodes is defined by a stochastic matrix **B** and a node assignment matrix **Z**; this model allows us to simulate different network structures by simply varying **B**, **Z** and *η*.

With this model in place, we first simulated networks with multiple non overlapping assortative communities of approximately the same size, in order to assess whether our model is able to recover known communities. In this case, **B** is a diagonal matrix with **B**_*ii*_ = *p*_SBM_, while the noise is added as an Erdos-Renyi network with a constant probability of connection *p*_ER_ (see Supplementary Materials); with these parameters, we then generated *k* blocks harbouring approximately *n/k* nodes each.

We then used the Signal to Noise Ratio (SNR) as an indicator of community detectability, such that we can measure whether the SBM model is distinguishable from background noise. The SNR is defined as follows:

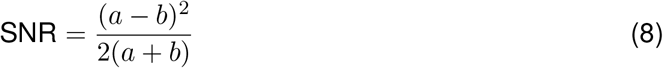

where *a* and *b* are the average degree within and outside the community (*a* = *np*_SBM_ and *b* = *np*_ER_) and *n* is the total number of nodes; by modulating *p*_ER_ and *p*_SBM_, we can control the SNR for a given network. Theoretically, SNR > 1 is the threshold for detectability, but it has been already shown that controlling for SNR > 1.5 is more reasonable [21].

However, since SBM-GNN is designed to detect multiple communities of different size, we simulated networks with hierarchical structure as follows; given a depth value, *h*, we generated *h* hierarchical layers of 2^*h*^ blocks each. Then, **B** is obtained as the average of the *h* layers, such that the smallest communities are the most assortative ones. Although there is no need to plant a hierarchical structure, this is a reasonable and realistic procedure to generate a network with multiple structured communities.

#### 2.2.2 Synthetic gene features

We have also generated community-aware features conditioned on community structure, such that nodes belonging to closer communities are more likely to share similar features. To do that, we generate correlated random variables using the feature coloring method, which is the inverse of features whitening, a method routinely used to remove correlation between random variables.

Specifically, we first generated features as independent random variables drawn from a Normal distribution, 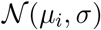, where the average *μ_i_* depends on the *i*-th community the node belongs to (see Supplementary Materials); then, the coloring procedure is applied such that features are conditioned on the SBM structure of the network, which leads to features that are probabilistically more similar for nodes within the same community (see Supplementary Materials).

#### 2.2.3 Community detection metrics

We then introduced two different metrics to assess community detection performances, namely the Jaccard coefficients, *J_c_*, and the assignment penalty.

In our case, we measured *J_c_* between each simulated block, *R_i_*, and each block learnt by SBM-GNN, 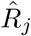, obtained by assigning the nodes to the block with highest membership probability. Since the node assignment to the blocks is order invariant, for each learnt block we used 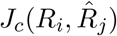, where 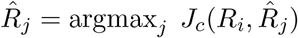.

However, Jaccard coefficients do not measure the uncertainty of the node membership. Thus, we defined the assignment penalty metric, *P_ij_*, between a known block assignment **Z**_:*i*_, that is the known assignment for each node to the *i*-th block, and the SBM-GNN assignment 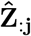, as 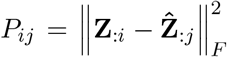, where 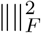 is the Froebenius norm. Similar to the Jaccard coefficient, we used the penalty metric for the j-th block as *P_j_* = min_*i*_(*P_ij_*). We then aggregated penalty scores into a single term, *P_tot_*, by summing all penalties and normalising them w.r.t the number of nodes and the number of blocks, such that 0 ≤ *P_tot_* ≤ 1.

### 2.3 Cancer genome data and protein interaction datasets

We downloaded genomic data from the Pan Cancer Analysis of Whole Genomes (PCAWG) project [22], used in the companion pathway and network analyses [23, 24]. We then obtained p-values associated with the confidence that a genomic locus is a driver; specifically, for each gene, we considered p-values for coding (CDS) regions and 4 different types of non-coding regions, namely 3’ UTR, 5’ UTR, promoters and enhancers, and then used Fisher’s method to transform p-values into *χ*^2^ statistics.

We considered five different cancer panels as gene labels, both for training our model and evaluating its performances, including the pathway implicated drivers (PID) gene list and the COSMIC Cancer Gene Census (Cosmic) [25] (see Table 3).

Finally, we used two protein-protein interaction datasets: the STRING [26] and BioGRID [27] database. We processed both datasets to keep only high confidence interactions [24] and connected components; here we found BioGRID to be smaller and less dense than STRING (see Table 2).

**Table 2:** Protein-protein interaction network properties. For each network, we report the number of nodes, the number of edges, the density, the maximum and median node degree, and the median clustering coefficient.

|  | # nodes | # edges | Density | Max degree | Median degree | Mean clustering |
| --- | --- | --- | --- | --- | --- | --- |
| STRING | 10224 | 205422 | 0.003931 | 1882 | 11.0 | 0.46084 |
| BioGRID | 5440 | 16346 | 0.001105 | 248 | 3.0 | 0.187394 |

**Table 3:** Cancer driver genes panels. For each cancer driver gene panel, we report the number of genes, and the number of genes mapped to genes in BIOGRID and STRING.

|  | # genes | BioGRID | STRING |
| --- | --- | --- | --- |
| PID | 169 | 134 | 161 |
| COSMIC | 719 | 481 | 610 |

## 3 Results

### 3.1 Performance on simulated data

We rigorously tested the performance of SBM-GNN on simulated networks generated by SBMs with controllable parameters, which is key to prove that our model is able to detect communities.

To do that, we simulated SBM networks with two blocks by drawing *p*_SBM_ ~ Uniform(0.5, 0.7) and adjusting the diagonal values of **B** to control the Signal to Noise Ratio (SNR), while using the identity matrix as gene features. Consistent with previously reported estimates, SBM-GNN was able to correctly detect the planted communities for SNR > 1.5 (see Figure 2A).

**Figure 1:**
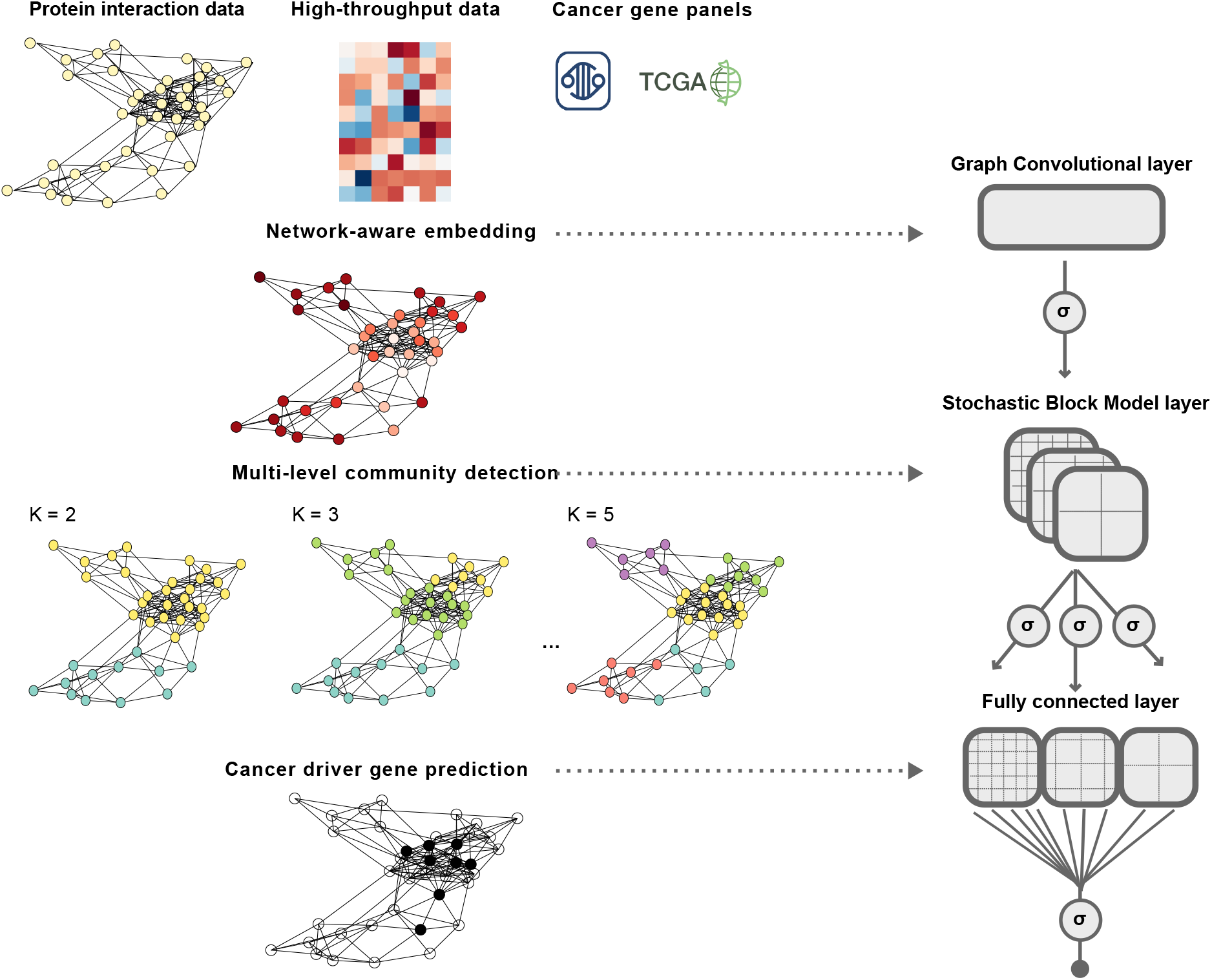
An overview of the Stochastic Block Model Graph Neural Network (SBM-GNN) model. Here we present a sketch of the processing steps (left) implemented by SBM-GNN into a deep neural network (right). Our model takes in input protein interaction data, high-throughput omics data and a cancer driver gene panel, which classifies each node as being a driver or not. Then, a Graph Convolutional layer is used to generate a gene-level network-aware feature embeddings, which in turn are used to assign genes to communities learned by multiple Stochastic Block Model (SBM) layers with varying number of blocks. Finally, the network structure learned by SBM layers is used as input for a fully connected classifier to predict cancer driver genes.

**Figure 2:**
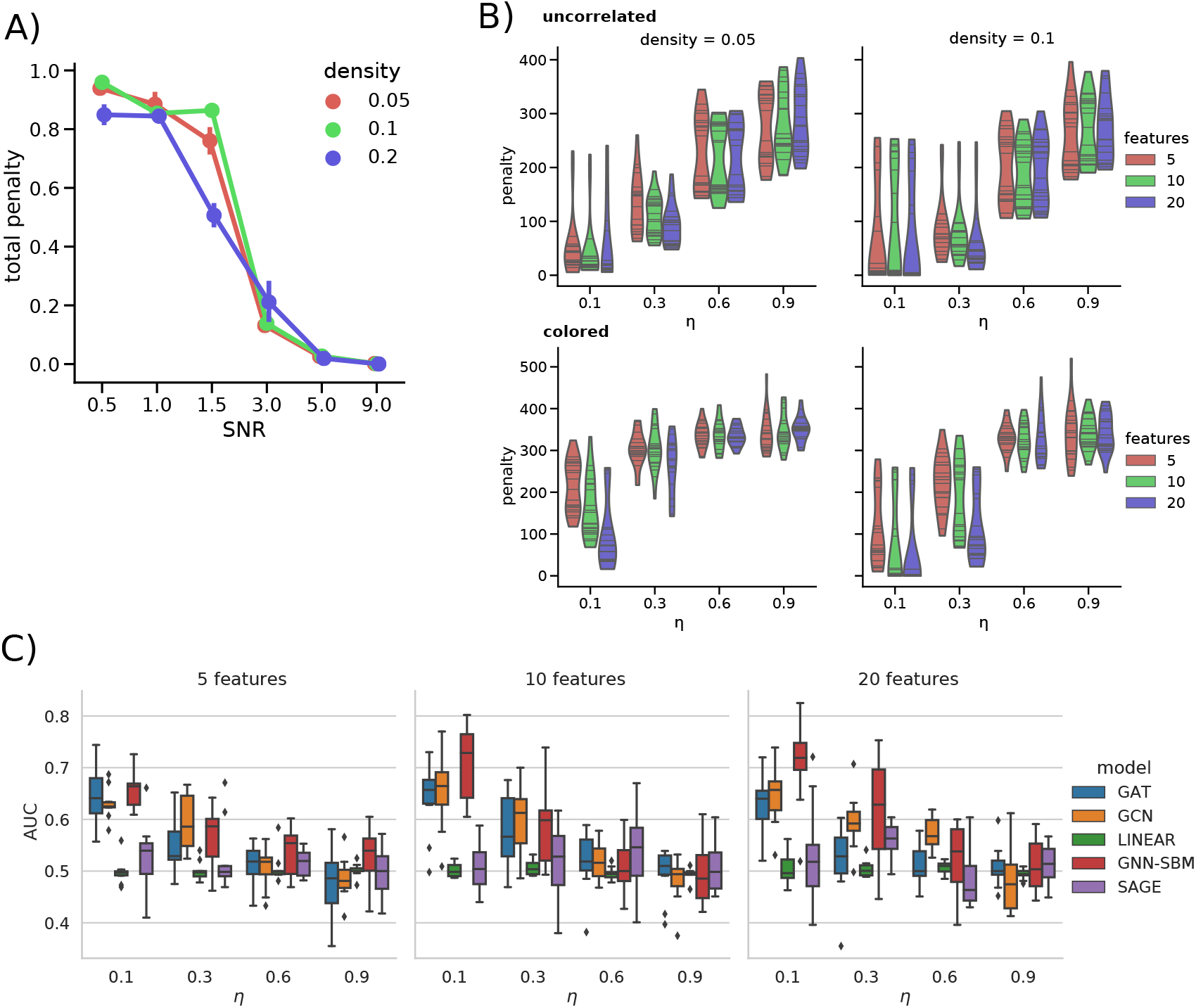
SBM-GNN performance on synthetic data. A) On the y-axis, we report the total normalised penalty obtained for 2 block SBM networks, whereas the x-axis reports SNR values and colors denote different density parameters. For for SNR= 0.5, non-detectable blocks, the total penalty is approximately 1, which corresponds to the maximum error. Total penalty drops for SNR> 1.5, confirming that our model is able to recover the network structure. B) Performance on uncorrelated and colored features for simulated networks with 4 blocks. On the x-axis, we show the penalty value on block assignments at varying levels of noise, *η*, where communities should become detectable for *η* < 0.5. We report results for 5, 10, 20 gene features (colored) and for network density *d* = {0.05, 0.1} (columns). C) Classification performance (AUC) of SBM-GNN, GAT, GCN,SAGE, and LINEAR architectures (color) on colored features and synthetic networks with 4 blocks. Results are shown for different *η* (x-axis) and for 5, 10, 20 features (columns). For low *η* values, all graph neural networks have performances significantly better than a random classifier (AUC> 0.5). Interestingly, as the number of correlated features increases, we observed a significant improvement on SBM-GNN performance.

We then assessed the performance of our method on networks with more than 2 communities and in the presence of gene features. In this case, we simulated networks of 1000 nodes with 4 blocks as a mixture of SBM structures, with noise weighted by *η* = {0.1, 0.3, 0.6, 0.9}. For each network, we then simulated genes with 5, 10, 20 features and generated 5 replicates for each possible parameters setting; in this case, we used both uncorrelated and colored features. Here we found that SBM-GNN was able to accurately detect communities in presence of more than 2 blocks and with multiple annotations (see Figure 2B). Moreover, by increasing the number of correlated features, which corresponds to strengthening the signal, performances clearly improved (see Figure 2B).

After confirming that SBM-GNN can detect communities, we tested its performances on node labelling and compared it to other GNN models, including GCN[10], GAT[13], SAGE[12], and a LINEAR model, that is a neural network made of two fully connected layers that ignores graph structure. While other, more complex, methods might have better performances on specific datasets, these three methods are usually at the foundation of most state-of-the-art approaches. We used our simulation framework to build networks and datasets with 5, 10,20 colored genes features and 10% of positive labels (see Figure 2C), ultimately generating 5 datasets for each possible parameters setting. For *η* < 0.5, signal is weighted more than noise, and all graph-aware methods yield better performances than a random classifier, whereas the LINEAR model always had Receiver Operating Characteristic (ROC) Area Under the Curve (AUC) *AUC* ~ 0.5. Interestingly, as the number of correlated features increases, SBM-GNN clearly outperformed the other models, suggesting that our approach is able to better exploits high-dimensional data.

Taken together, our simulations showed that SBM-GNN is a robust and effective architecture to identify network communities and to predict node labels using high-dimensional gene-level information.

### 3.2 Performance on pan-cancer genome data

We then used our model to perform cancer driver gene prediction and cancer pathways discovery using pan-cancer genomic data [3] and two protein-protein interaction datasets, namely STRING and BioGRID, while using the PID gene panel as node labels for training. Here we used both our base architecture and a set of 9 other SBM-GNN extensions, since fine tuning the layers and parameters is often key to yield the best performances in deep learning (see Supplementary Materials).

Interestingly, we found that all architectures performed remarkably well (AUC > 0.7) regardless of the protein interaction dataset used, albeit STRING seems to provide consistently better results (see Figure 3A). Moreover, we found that adding the residual information consistently improved the performances of our model, and similarly, but to a lesser extent, the use of graph diffusion.

**Figure 3:**
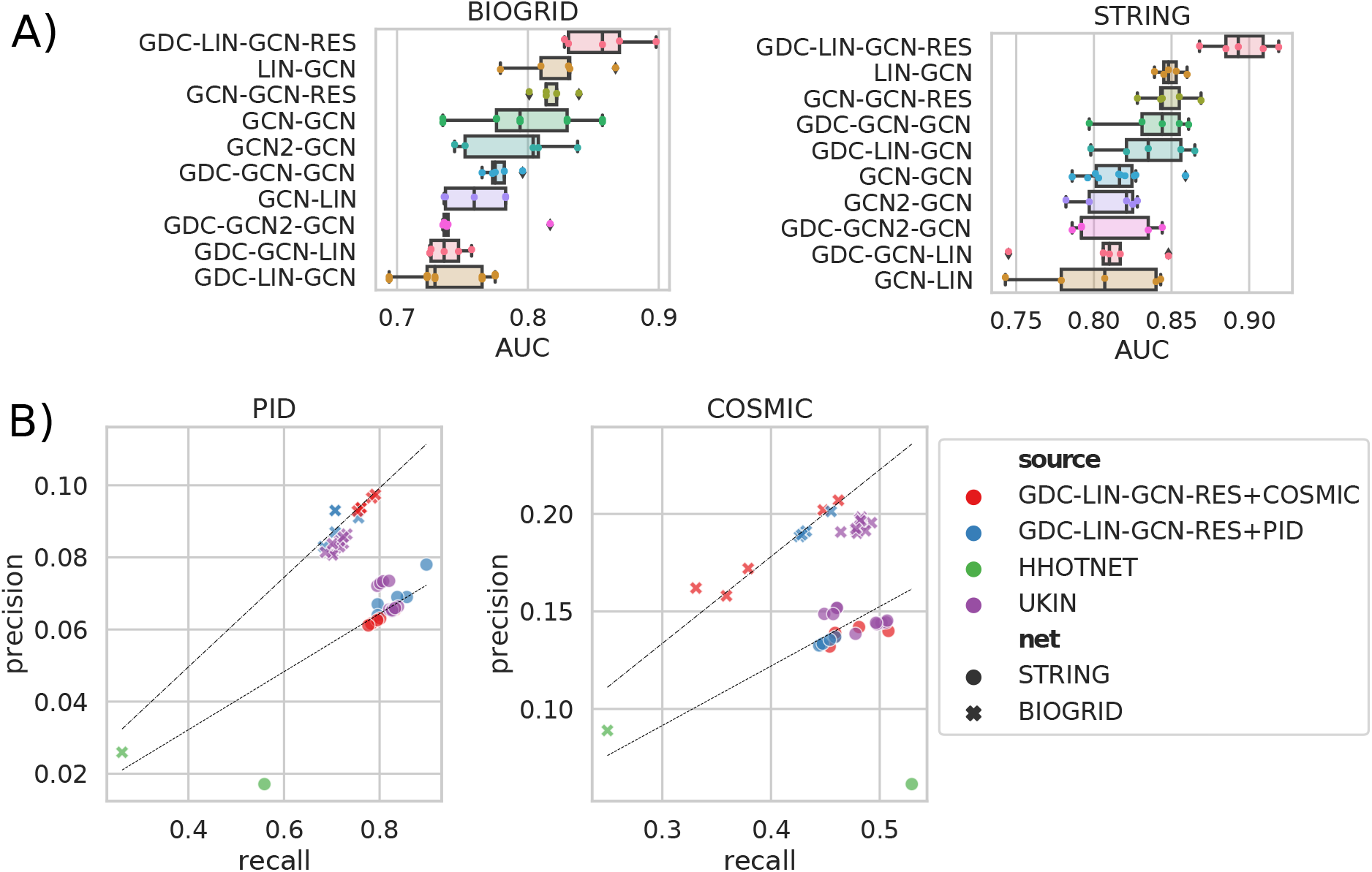
SBM-GNN performance on cancer driver genes prediction. A) Cancer driver gene prediction measured as the area under the receiver operating characteristic curve (ROC AUC), for different SBM-GNN architectures. We have sorted them by average performance over 5 runs. For both the BIOGRID and STRING networks, we found GDC-LIN-GCN-RES architecture achieves the best performances; interestingly, adding residuals information increase the AUC for both SGCN-GCN and LIN-GCN models. It is also worth noting that all neural networks without a network-aware block assignment layer, namely GCN-LIN, GDC-GCN-LIN, have consistently worse performances. B) We compare the recall (x-axis) and precision (y-axis) of hierarchical hotnet (HHotNet), UKIN, and SBM-GNN (colors) on BIOGRID and STRING networks (markers). Dashed lines are the theoretical relationship between precision and recall at the specific percentile for each network. For UKIN and SBM-GNN, we reported as cancer driver genes those above the 80th percentile of their scores. Here, we found SBM-GNN with residuals to be the best performing method, a trend we also observed when we trained our model on the COSMIC gene panel.

We then compared the performance of SBM-GNN with that of GCN [10], GAT [13], SAGE [12], and a basic model with two fully connected layers, LINEAR (see Supplementary Materials). Most SBM models achieved comparable performances with the other deep learning models (see Table 4), but the residual version of SBM-GNN consistently achieved the best performances, with GCN being the best alternative.

**Table 4:** Performance analysis of different models on cancer driver genes prediction. Models were trained using either the BIOGRID or the STRING protein-interaction network, and the PID panel of cancer driver genes. For each model, we report the precision, recall, balanced accuracy (BACC) and Area Under the ROC Curve (AUC) averaged over 5 independent runs. We report in bold face the best performance for each metric, and the best overall model.

| Network | Model | Precision | Recall | BACC | AUC |
| --- | --- | --- | --- | --- | --- |
| BioGRID | GAT | 0.0798 | 0.6194 | 0.7174 | 0.8086 |
|  | GCN | 0.0798 | 0.6294 | 0.7212 | 0.8282 |
|  | LINEAR | 0.0250 | **1.0000** | 0.5000 | 0.7814 |
|  | SAGE | 0.0832 | 0.7024 | 0.7516 | 0.8236 |
|  | GDC-LIN-GCN | 0.0662 | 0.5316 | 0.6684 | 0.7372 |
|  | **GDC-LIN-GCN-RES** | **0.0894** | 0.7120 | **0.7626** | **0.8568** |
|  | GCN-GCN | 0.0764 | 0.5952 | 0.7046 | 0.7984 |
|  | GCN-GCN-RES | 0.0772 | 0.6440 | 0.7226 | 0.8180 |
| STRING | GAT | 0.0418 | 0.8366 | 0.6620 | 0.6844 |
|  | GCN | 0.0604 | 0.7554 | 0.7824 | 0.8660 |
|  | LINEAR | 0.0210 | **0.9346** | 0.5420 | 0.8150 |
|  | SAGE | 0.0630 | 0.8044 | 0.8052 | 0.8626 |
|  | GDC-LIN-GCN | 0.0524 | 0.6654 | 0.7358 | 0.8350 |
|  | **GDC-LIN-GCN-RES** | **0.0694** | 0.8368 | **0.8270** | **0.8948** |
|  | GCN-GCN | 0.0508 | 0.6368 | 0.7218 | 0.8046 |
|  | GCN-GCN-RES | 0.0598 | 0.7554 | 0.7810 | 0.8478 |

#### 3.2.1 Comparison with state-of-the-art cancer driver prediction methods

We then compared SBM-GNN performance with state-of-the-art methods for cancer driver gene prediction (see Supplementary Materials). There is a vast literature on methods designed to identify new cancer driver genes, with many of them using network information [8, 28, 29]. Recently, the *using Knowledge In Networks* (uKIN) [8] has been shown to be the gold standard in the field, and hence we used it as a benchmark to analyse SBM-GNN performance. Conversely, methods for community detection are less established. There are a plethora of methods that detect cancer-associated submodules, that are connected subset of genes driving cancer [9, 30], but they do not explain the structure of all the nodes that are not implicated in cancer. However, for completeness, we compared SBM-GNN performance with Hierarchical Hotnet (HHotNet) [9] to explore the performance of this class of methods (see Supplementary Materials). It is worth noting that HHotNet and uKIN are unsupervised methods, albeit the latter uses information on known cancer driver genes to guide the diffusion process. For this reason, we carried out comparisons both with models trained and tested on the PID labels, and by training on the COSMIC panel and testing on PID. SBM-GNN performance was consistent with those of uKIN and outperformed it when using PID labels (see Figure 3C). Unsurprisingly HHotNet had a clearly worse performance. While results are not directly comparable, it is interesting to observe that submodule inference is not directly applicable to identify new cancer drivers.

#### 3.2.2 Discovering cancer pathways by inspecting stochastic block models

Our method is not limited to the prediction of cancer driver genes, but more importantly provides an interpretable picture linking cancer driver genes to the pathways they are involved in. Our hypothesis is that cancer driver genes are targeting multiple, possibly distant pathways, and that complex relationships might be captured by fitting multiple SBMs. Thus, using our model, we analysed the blocks harbouring cancer driver genes, the relationship between SBMs, and how the learnt blocks can be used to identify cancer pathways.

We selected the model with the lowest loss and analysed the genes assigned to each block. As we hypothesised, cancer drivers genes are assigned to multiple blocks, both in the coarser and finer SBM layers (see Figure 4). This observation is consistent with the fact that cancer driver genes, such as *TP53* or *MYC*, are involved in multiple biological processes. Interestingly, in most cases, cancer driver genes represent less than 20% of the genes in a block, which is consistent with a model of tumorigenesis where driver genes mediate cancer phenotypes by interacting with non driver ones.

**Figure 4:**
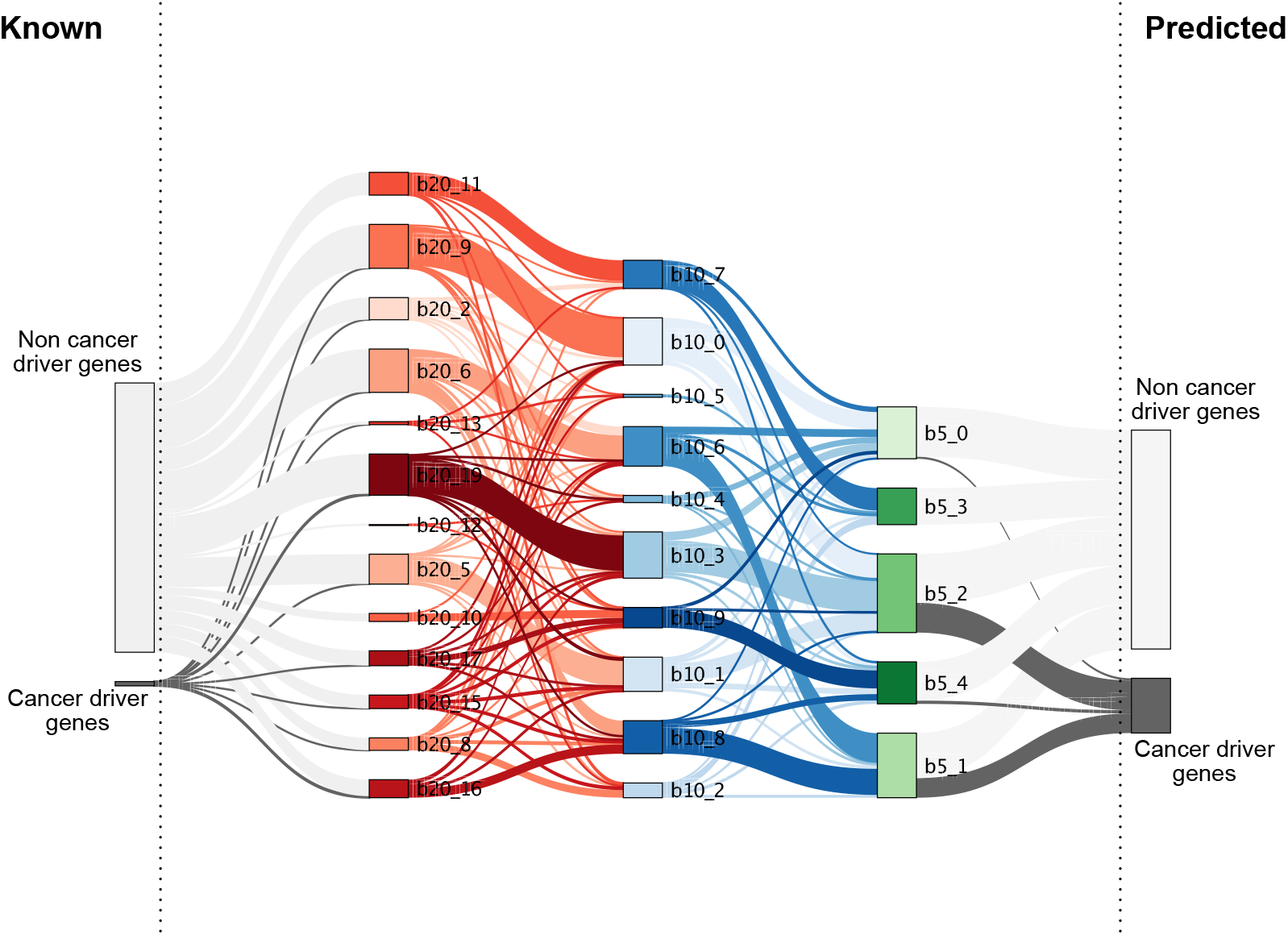
Block level organization of cancer driver genes. From left to right, we show the cancer labels, the blocks of size *k_s_* = {20, 10, 5} and the predicted labels. Between different nodes, we are plotting branches representing the number of nodes shared between the two blocks. The last blocks on the right are those that SBM-GNN predicts as significant.

Finally, we functionally characterised the genes in the blocks identified by SBM-GNN, in order to provide a system-level picture of gene organisation encoded by the SBM. To do that, for each block of genes, we performed a Fisher’s exact test using the Reactome genesets (see Figure 5). Since we are organising the network in communities, we expect to find blocks recapitulating cancer-associated pathways, alongside others not mediating cancer phenotypes.

**Figure 5:**
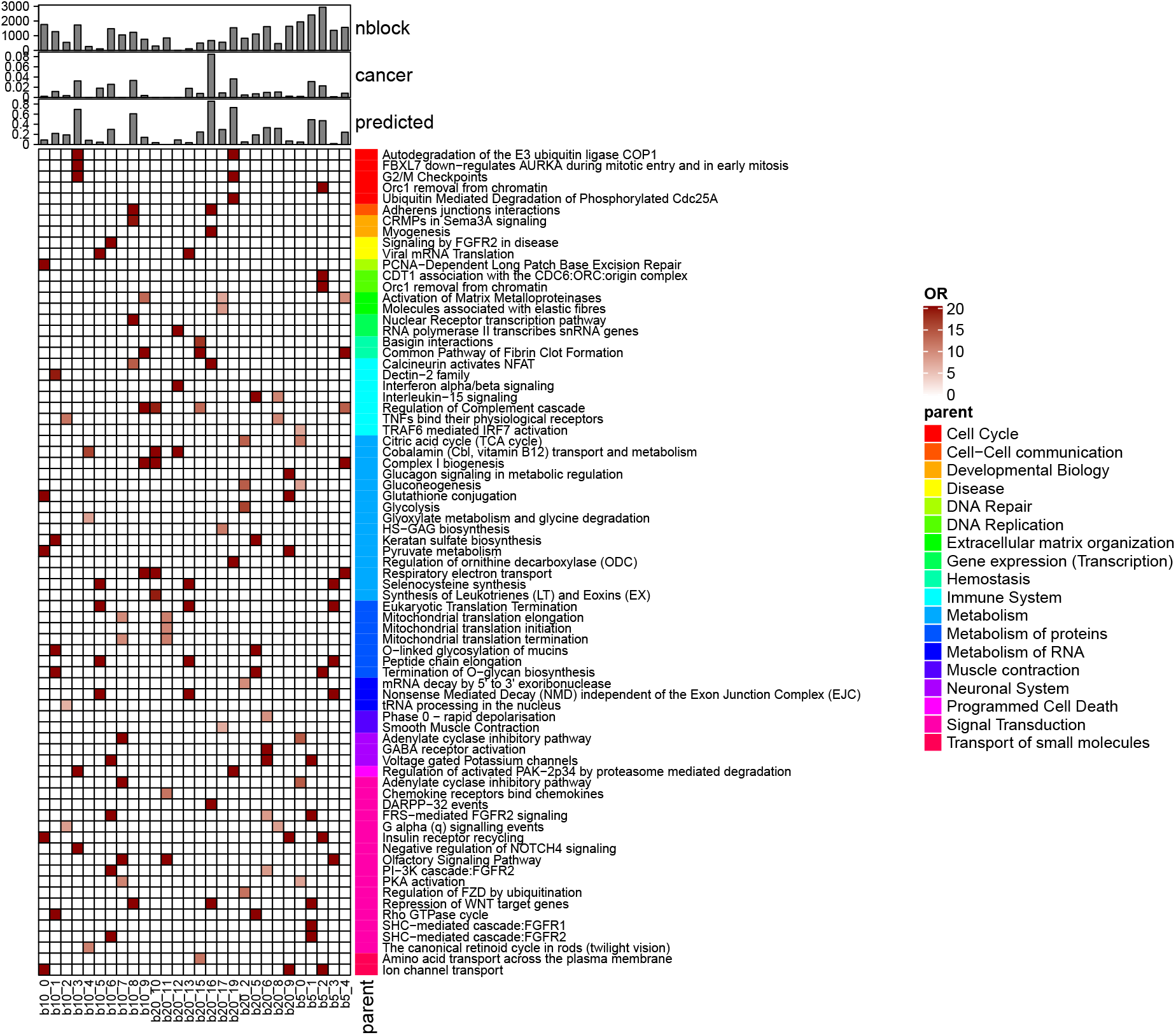
Block level Reactome geneset enrichment analysis. For each block (columns), we plot the top five statistically significant genesets ranked by odds-ratio (OR) from the Fisher’s exact test. For each Reactome geneset (rows), we also report their parental group (color annotation), which allows us to identify macro functional classes. For each block, we show, from top to bottom, the number of genes in the block, the number of known cancer driver genes in the block and the fraction of those predicted by our model.

At the higher level, our model identified blocks of genes associated with hallmarks of cancer [31], in particular programmed cell death, metabolism which is associated with deregulation of cellular energetics, and genome instability through alteration of the DNA repair and replication machineries. Specifically, by looking at the 20 block SBM, we found one (b20 16) having 86% of its genes been predicted as cancer driver genes, which recapitulates 72% (OR:45.1, p: 1.43 × 10^−8^) of genes involved in the *Calcineurin activates NFAT* process, that is a T-cell related process involved in cancer progression and metastasis [32, 33], 70% (OR: 44.4, p: 2.32×10^−22^) of the genes involved in Adjerens Junctions Interactions, and 60% (OR: 23.6, p: 1.11 × 10^−6^) of the genes associated with *Repression of WNT target genes* pathway, which are both well known processes involved in cell motility and proliferation. Moreover, we also found another block (*b*20 6) encompassing the PIK3 cascade pathway (OR: 5.74, p: 2.07 × 10^−6^) and multiple *FGFR1* and *FGFR2* signalling processes, which are known to be critical for tumorigenesis [34]. Conversely, for example, block B20 11, which does not harbor any cancer driver gene, is associated with non-cancer related processes, such as *Olfactory Signaling*.

Taken together, we have shown that our model is able to recover genes and pathways associated with well characterised cancer mediating processes; in particular, by learning SBMs with an exponentially growing number of clusters, it is possible to go from broad molecular hallmarks to specific biological processes.

## 4 Conclusions

Tumorigenesis is triggered by a complex molecular reprogramming mediated by genetic, genomic, and molecular alterations acquired by driver and non-driver genes. While it is now possible to quantitatively assess the impact of these changes at the single gene level, reconstructing a system-level picture of cancer cells remains a challenging task.

Here we introduced a new model, called SBM-GNN, combining recent geometric deep learning architectures with stochastic block model to simultaneously infer cancer driver genes and associated pathways; our model provides a scalable approach to integrate multi-omic data with protein-interaction information, that can be used to generate testable hypothesis at the gene and pathway level. We validated our method using an extensive set of simulations showing that SBM-GNN can correctly identify cancer driver genes and cancer related pathways.

We then applied our method to the analysis of pan-cancer genomic data, where we showed that SBM-GNN can predict driver genes with high accuracy and identify blocks of genes associated with well-know hallmarks of cancer. On this point, the ability of our model to learn an easily interpretable pathway organization of cancer genes provides new opportunities to dissect the system-level reprogramming underwent by cancer cells.

## Contributions

G.S. and V.F. conceived the study. V.F., G.S. and P.L. designed the model. G.S., V.F. P.L. and R.V.T. developed the simulation framework. V.F. wrote the software and performed all analyses, supervised by G.S. G.S., V.F. and P.L. analysed the data. G.S. and V.F. wrote the manuscript with contributions from all the authors.

## Acknowledgements

This work has been supported by the Wellcome Trust Seed Award in Science (207769/A/17/Z) to G.S. The GPU hardware used for this research was donated by the NVIDIA Corporation to G.S.

