## Supplementary Materials for "Discovering cancer driver genes and pathways using stochastic block model graph neural networks"

### 1 Methods

#### 1.1 Simulated features

In our simulations, we studied the performance of our model using both uncorrelated and correlated gene features.

In the first case, we simulated features conditioned on the communities genes belongs to, but we did not account for correlation between different communities. Given a planted SBM structure, we simulated features for each community as random samples from a normal distribution; in the case of a hierarchical SBM, we only considered the blocks in the deepest layer. Specifically, given $K$ communities, we first selected at random their mean from a uniform distribution $\mu_k$ = Uniform$(-5, 5)$, and we then sampled the features for each gene, $i \in k$, as $\mathbf{X}[i,:] = \mathcal{N}(\mu_k, 1)$. After obtaining the feature matrix $\mathbf{X}$, we set a fraction of genes $N_{cancer}$ as cancer genes and multiplied their features by a weight factor $W$. With this strategy, features have probabilistically a distinct signal from the background.

We then used the Cholesky decomposition method to generate correlated features. This process, called features coloring, allows to impose a covariance matrix onto a stochastic process such that the final samples are correlated. Given $\mathbf{Y}$ a random variable of i.i.d. noise (uncorrelated) and $\mathbf{\Sigma}$ a covariance matrix, we want to find $\mathbf{X}$ s.t. its values are correlated conditioned on $\mathbf{\Sigma}$. Given a positive, semi-definite matrix $\mathbf{\Sigma}$, the Cholesky decomposition finds a lower triangular matrix $\mathbf{L}$ $s.t.$ $\mathbf{\Sigma} = \mathbf{L}\mathbf{L}^T$. Correlated samples can then be found as:

$$\mathbf{X} = \mathbf{L}\mathbf{Y} \tag{1}$$

However, this approach requires the adjacency matrix to be positive semi-definite, which is not often the case. To overcome this limitation, we used the covariance matrix $\mathbf{\Sigma} = \mathbf{A}\mathbf{A}^T$, which can be considered an edge correlation matrix.

#### 1.2 Performance metrics

##### 1.2.1 Blocks assignment and characterisation

One main advantage of the SBM-GNN model is the interpretability of the hidden layers. We remind that in the main manuscript we have described (Eq. 2-3) how the the membership matrix $\mathbf{Z}^{(s)}$ for the $s$-th SBM is learnt and then concatenated into the last layer. The block assignment matrix $\mathbf{Z}^{(s)}$, can be used to understand how the genes are assigned to different blocks, the relationship between different blocks and their characteristics.

Given the matrix $\mathbf{Z} = \mathbf{Z}^{(1)}|\mathbf{Z}^{(2)}\ldots|\mathbf{Z}^{(S)}$, each sub-matrix $\mathbf{Z}^{(s)}$ has dimensions $n \times k_s$, with $n$ being the number of nodes and $k_s$ the number of blocks in the $s$-th SBM, respectively. Each row represents the probability of the node to belong to one of the $k_s$ blocks (soft assignment). The $\mathbf{Z}$ matrix is saved alongside the neural network parameters and can then be used to assess the community assignment performance with simulated data, when the background blocks are known, or to characterise the blocks obtained with cancer data.

Here we use a toy example to illustrate how $\mathbf{Z}$ is used in practive. Given an architecture with 2 parallel groups of respectively 2 and 4 blocks, the resulting matrix $\mathbf{Z}$ would be of the form:

| | **b2_0** | **b2_1** | **b4_0** | **b4_1** | **b4_2** | **b4_3** |
|---|---|---|---|---|---|---|
| **gA** | 0.1 | **0.9** | 0 | 0.3 | **0.5** | 0.2 |
| **gB** | 0.2 | **0.8** | 0 | **0.9** | 0.1 | 0 |
| **gC** | 0.3 | **0.7** | 0 | 0.3 | **0.5** | 0.2 |
| **gC** | **0.8** | 0.2 | 0 | **0.9** | 0.1 | 0 |
| **gD** | **0.9** | 0.2 | 0 | 0.3 | **0.6** | 0.1 |
| **gE** | 0.4 | **0.6** | 0 | **0.6** | 0.4 | 0 |
| **gF** | **0.6** | 0.4 | 0 | 0.1 | **0.9** | 0 |
| **gG** | **0.7** | 0.3 | 0 | 0.3 | **0.5** | 0.2 |
| **gH** | 0.1 | **0.9** | 0 | **0.9** | 0.1 | 0 |

where each entry is the probability that the node, rows gA,gB ... gH, belongs to one of the blocks, b2_0, b2_1, ..., b4_3. Here the block naming convention, e.g. b2_1, indicates both the group (SBM with two blocks ) and the specific block within the group (second block). The softmax function is applied for each group (b2 or b4) such that all nodes are assigned to each parallel SBM.

Eventually, nodes are uniquely assigned to a single block for each group by picking the block with the largest memebership probability. In this example, we would obtain the assignment :

- b2_0: gC, gD, gF, gG
- b2_1: gA, gB, gE, gH ...

#### 1.3 SBM-GNN hyperparameters

We trained our model with the following hyperparameters: learning rate:0.01, weight decay:1e-4, 16 hidden nodes (for the $\phi$ hidden layers), and 3 parallel SBM with 5,10,20 blocks (hidden nodes of $\zeta$). Training was done over 15,000 epochs, with 80% of nodes in the training set and 10% in the test set. The same parameters were used with the simulated data, however we trained the model on 1,000 epochs, as they were sufficient to train SBM-GNN on a network with only 1000 genes.

#### 1.4 UKIN and Hierarchical HotNet

We compared the performances of our model with those of two other state-of-the-art network-aware analysis methods: *using Knowledge In Networks* (uKIN) and *Hierarchical HotNet* (HHotNet).

uKIN and HHotNet are network inference methods for attributed networks, although they allow only single gene features; thus, we used Fisher's method to combine multiple datasets from PCAWG into a single score. uKIN also requires a set of known cancer driver genes to act as seeds for the guided random walks; similar to what proposed by the authors of the method, for each run (10 in total), we randomly sampled 30 cancer genes from the COSMIC dataset and used them as seeds. Testing is then done with the whole COSMIC geneset, but those employed for training, and with the PID labels, which are those deemed as cancer genes specific to this dataset. Results are presented in Fig. 5.

Conversely, Hierarchical Hotnet is a method to detect disrupted cancer subnetworks. It is then important to notice that the overall classification performance is probably worse than the one of UKIN as it is not directly designed to extract single cancer drivers but submodules of them. We ran Hierarchical Hotnet with the score randomisation strategy, 100 permutations, and extracted all the modules returned by it.

#### 2 Figures

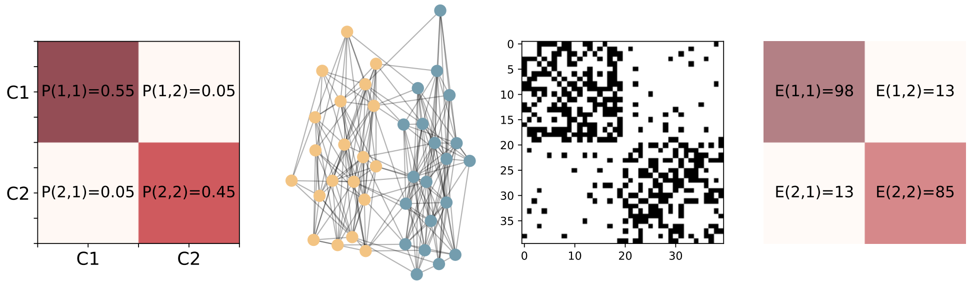


Figure 1: **Stochastic Block Model, network parameters and generation.** Here we present a simple stochastic block model of a network with two communities. We generate a network with $40$ nodes, with $20$ nodes assigned to community $C_1$, and the other $20$ nodes to $C_2$. The probability of connection between nodes is defined by the SBM matrix shown on the right. Each element of the matrix defines the probability of connection within and between the blocks. Next to the community matrix, from left to right, we show a network generated from our SBM, the corresponding adjacency matrix, the number of observed links between each block on the left. In the network, different colors denote nodes in different communities, where the adjacency matrix is sorted by blocks. First, we can notice that a higher probability leads to more edges between the nodes; in this case we have an assortative network, where nodes within the same community are more connected than nodes between different communities.

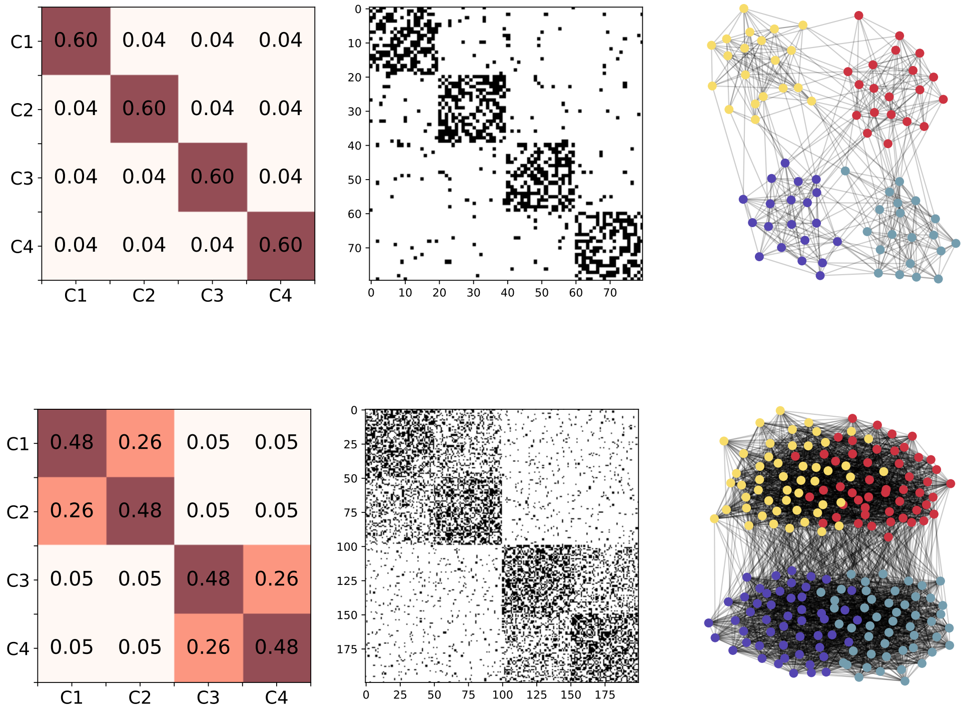


Figure 2: **SBM matrices for both disjoint and hierarchical communities** We show here two examples of how we simulate synthetic networks. First, top row, we simulate disjoint assortative communities with $p_{ii}$ between $0.5$ and $0.7$ (values on the diagonal) and we add some background noise, by setting $\eta = 0.1$. On the right side we show a network generated from our SBM, with different colors to denote different communities, and the corresponding adjacency matrix. We then show a hierarchical SBM community matrix (bottom row). Blocks C1 and C2, and C3 and C4, are merged together to generate a hierarchical structure in the network. Connection probability $P_{i,j}$ is averaged across hierarchical levels.

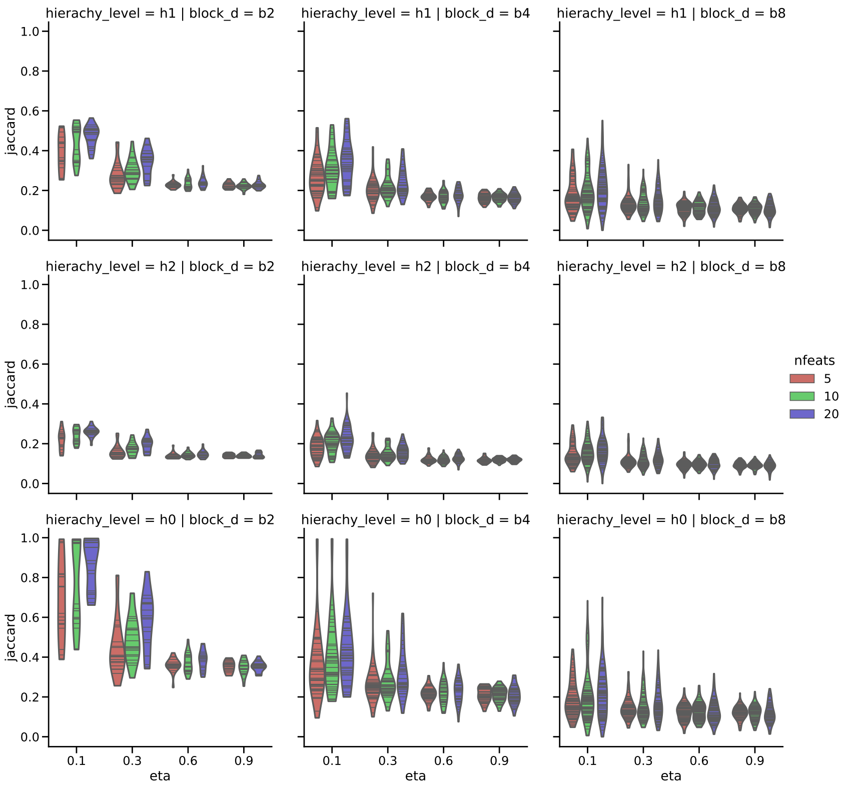


Figure 3: **Jaccard coefficient of colored features and 8 blocks.**Max Jaccard for each simulated dataset and run, for different values of $\eta$ (x-axis), different number of features (color), hierarchy level (columns), block level (rows). We can notice that the detection for 2 blocks (h0 and blue dots) is good. For 4 and 8 blocks instead, it seems that SBM-GNN is not able to fully recover the fine structure, leaving some blocks empty, but properly recognising the others. We can also notice good performance for low noise (eta = 1) while expected randomness for random networks (eta = 0.9)

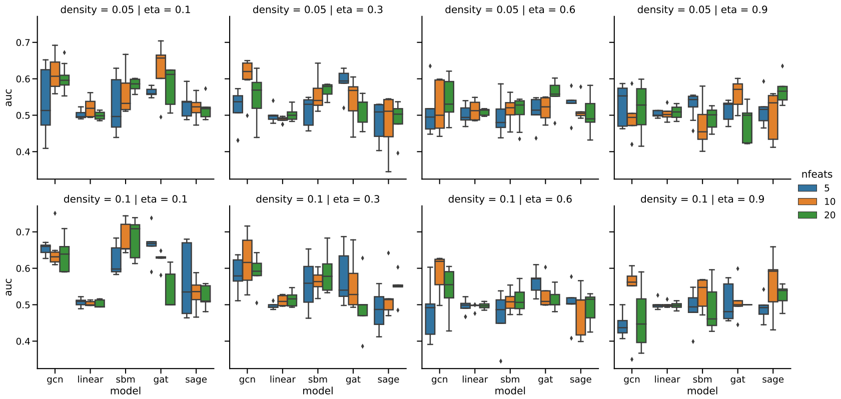


Figure 4: **Classification of colored features and 8 blocks.**AUC performance of different architectures (x-axis) for different $\eta$ (columns), density (rows), number of features (colors).

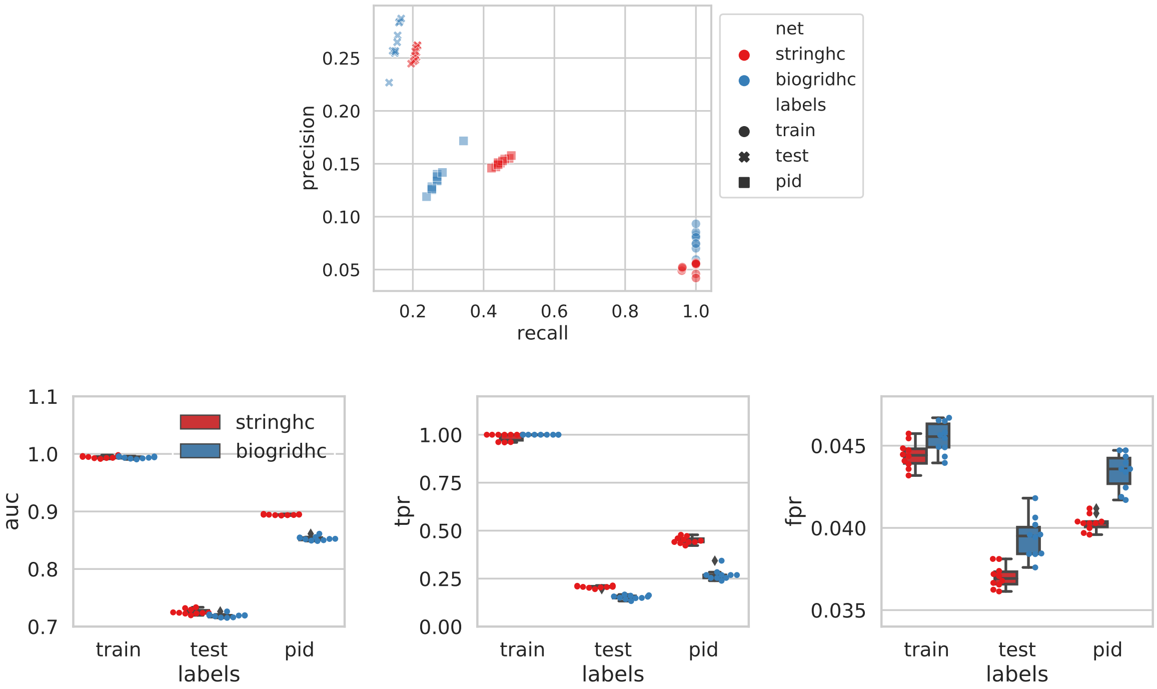


Figure 5: **UKIN performance**. A) Precision and recall for uKIN using different networks (colors) and on different labels, we cosider as significant the genes above the 90-th percentile. Train performance (round marker) is for the same genes used as seeds. The test performance (crosses) is that on the cosmic geneset, excluding the seed genes. The PID performance is on the original cancer genes of the dataset (square marker). B) AUC, TPR and FPR for the same data shown before.
